# Base pairing and stacking contributions to double stranded DNA formation

**DOI:** 10.1101/2020.08.22.262667

**Authors:** Martin Zacharias

## Abstract

Double-strand (ds)DNA formation and dissociation are of fundamental biological importance. The negatively DNA charge influences the dsDNA stability. However, the base pairing and the stacking between neighboring bases are responsible for the sequence dependent stability of dsDNA. The stability of a dsDNA molecule can be estimated from empirical nearest-neighbor models based on contributions assigned to base pair steps along the DNA and additional parameters due to DNA termini. In efforts to separate contributions it has been concluded that base-stacking dominates dsDNA stability whereas base-pairing contributes negligibly. Using a different model for dsDNA formation we re-analyze dsDNA stability contributions and conclude that base stacking contributes already at the level of separate ssDNAs but that pairing contributions drive the dsDNA formation. The theoretical model also predicts that stability contributions of base pair steps that contain only guanine/cytosine, mixed steps and steps with only adenine/thymine follows the order 6:5:4, respectively, as expected based on the formed hydrogen bonds. The model is fully consistent with available stacking data and nearest-neighbor dsDNA parameters. It allows to assign a narrowly distributed value for the effective free energy contribution per formed hydrogen bond during dsDNA formation of −0.72 kcal·mol^-1^ based entirely on experimental data.

## Introduction

Complementary single-stranded (ss)DNA molecules can associate to form a DNA double helix (dsDNA) involving hydrogen bonds between A:T and G:C base pairs and stacking interactions between adjacent base pairs ^1,2^. It is well known that the stability of the DNA double helix depends on the base composition ^3–9^. For example, it has been found that the stability of long DNA double strands of random sequence depends linearly on the G:C content^2,8,10^. Nearest neighbor models have been developed to account for the sequence dependence of dsDNA stability^5^. These models are parametrized based on melting data of DNA polymers, DNA oligomers of various sequences and lengths and DNA dumbbells ^5,6,11–13^. Nearest neighbor models have been used quite successfully for calculating the stability and melting behavior of DNA. Similar nearest neighbor models have been developed for dsRNA and are widely used for secondary structure prediction of RNA ^14^ and for estimating the stability of folded RNA molecules ^15^. However, the nearest neighbor parameters extracted from experiments include the sum of base pairing and (nearest neighbor) base pair stacking to the dsDNA stability. A separation into stacking and base pairing contributions could be helpful to understand many other aspects of DNA conformation and changes in DNA conformation during processes such as DNA replication, DNA repair due to damage or strand separation due to kinking or over- and unwinding of the duplex^8^.

Several theoretical studies have addressed the base pairing and stacking geometry and stability of DNA base pairs. These include quantum mechanical approaches that typically are performed on one or a few optimized geometries^16–18^. Such calculations predict that both stacking and base pair hydrogen bonding contribute to double strand stability. However, the neglect of the aqueous environment in QM calculations may over stabilize the influence of polar interactions. Molecular dynamics simulations based on a molecular mechanic force field and including explicit solvent indicate that isolated DNA bases but also dinucleotides and also longer single strands tend to stack in aqueous solution depending on the bases or nucleotide sequence^19–24^.

Several experimental approaches have also been used to investigate stacking energetics in dinucleotides as well as isolated nucleo-bases ^4,25–28^. Studies on the influence of dangling terminal bases that do not form base pairs with the opposite DNA strand are particularly interesting because base pairing hydrogen bonds do not participate^5,29,30^. However, dangling effects are small and since the dangling base has no partner and is at the terminus of a duplex it is not clear if the stacking exerts the same effect as a stacked base pair in a full duplex molecule. However, it is possible to introduce nicks, that are breaks in the sugar phosphate backbone in one strand, in dsDNA to study the stacking energetics in fully duplex DNA^13,31–34^. The unstacked form of a nicked DNA runs much slower and separate from the stacked (near B-DNA form) duplex DNA in polyacrylamide gel electrophoresis ^32^. By varying the denaturation conditions in the gel it is possible to directly measure the equilibrium between the stacked and unstacked form and upon extrapolation to zero concentration of denaturant it is possible to extract the free energy difference between a fully stacked and unstacked state at the nicked site. In this way it has been possible to obtain a full set of stacking free energies at nicked sites for all possible base pair steps ^13,32^. The free energy of base pair stacking at nicked states has also been studied by MD free energy simulations resulting in a reasonable correlation to experiment and indicating that the stacked structure at the nick is indeed similar to the stacking in dsDNA ^31^. Alternatively, base pair stacking has also recently been studied using single molecule manipulation with optical tweezers in combination with the DNA origami technology on DNA blunt-end stacking ^35^.

Based on the assumption that the base pair stacking quantified in DNA nicking experiments indeed represents the stacking contribution to the stability of dsDNA Frank-Kamenetskii and coworkers partitioned the dsDNA stability parameters into base pairing and stacking contributions ^13,32^. This model suggests that the stacking contribution largely drives dsDNA formation and that base pairing makes little or even a slightly opposing contribution to dsDNA stability ^8^. In the present study evidence is provided that the stacking energetics extracted from nicking (or other) experiments can also be interpreted as the tendency to stack already in the unpaired ssDNA. The dsDNA formation is then not a process starting from fully unstacked DNA to a fully stacked and base paired duplex. Instead the base pairing drives the dsDNA formation and it needs even to be strong enough for driving residual unstacked DNA into a fully stacked state in the dsDNA. The model provides an explanation for the nearest neighbor parameters in terms of base pairing and stacking contributions. It also predicts that the base pairing contributions to the stability of base pair steps that contain only G or C, mixed steps and steps that only contain A or T follows approximately the order 6:5:4 (or 3:2.5:2), respectively, as one would expect from the number of formed hydrogen bonds. The implications of the model will be discussed.

## Materials and Methods

Umbrella Sampling free energy simulations on dsDNA formation/separation of dinucleotide steps were performed using the Amber18 simulation package ^36^. A standard B-DNA duplex formed by two (self)complementary DNA dinucleotides (ApT) was built using the NAB module of the Amber18 package. The system was solvated with TIP3P water ^37^ in an octahedral box with a minimum of 20 Å between the DNA and the box boundaries. The ion concentration was adjusted to ∼0.1 M with added Na+ and Cl-ions (neutral system). After energy minimization (2000 steps) the system was heated in steps of 100 K up to a temperature of 300 K keeping positional restraints on all non-hydrogen atoms with respect to the start structure. The positional restraints were removed within another 2 ns equilibration at 300 K and constant pressure of 1 bar.

Umbrella sampling was performed along a distance coordinate between the centers of mass of the non-hydrogen atoms of two nucleo-bases on opposite strands (see Supporting Information, Figure S1 and Table S1). The relative angular orientation of the first base of one strand and the opposite base (second base of the second strand) was restraint by angular and dihedral restraints summarized in Table S2. A quadratic potential was added to the force field to control the reaction coordinate with equilibrium distances changing from 6 to 14 Å (see Table S1). Sampling in each US window was performed for 50 ns (after a 5ns equilibration phase). Two sets of simulations were performed. In the set A, the bases in each dinucleotide were allowed to unstack during every stage of the US simulation. In a second set B, the unstacking in each dinucleotide was suppressed by adding an angular restraining term (Supporting Information, Table S2). Hence, in this case the dinucleotides remained stacked even after dsDNA dissociation. The free energy change (potential-of-mean force) along the reaction coordinate was evaluated using the weighted histogram analysis (WHAM) method ^38^.

## Results and Discussion

According to the nearest neighbor model for DNA stability the energetics of DNA duplex formation can be calculated from contributions of overlapping base pair steps and additional terms due to the DNA termini (Table 2). Hence, the stability of a short duplex segment embedded in a longer DNA, e.g of sequence 5’-GCAT (with complementary strand: 5’-ATGC) can be calculated by adding the nearest neighbor parameters of GC, CA and AT base pair steps. With current consensus (unified) nearest neighbor parameters it is not only possible to reasonably predict the stability of dsDNA but also to predict the linear dependence of the melting stability of long random DNA sequences on the G/C content. This is surprising because the equation to calculate the stability of random DNA sequences by Protozanova et al. (2006) ^32^ includes not only terms linear in the G/C content but also terms quadratic in the G/C content. However, the coefficient of the quadratic term is close to zero with the current nearest neighbor parameters.

**Table 1.** Nearest neighbor base pair and stacking parameters

| Base pair step<br>5'-3'/5'-3' | $\Delta G_{\text{KL,SantaLucia}}$<br>(kcal·mol<sup>-1</sup>) | $\Delta G_{\text{KL,nick}}$<br>(kcal·mol<sup>-1</sup>) | $\Delta G_{\text{KL,SantaLucia}} - \Delta G_{\text{KL,nick}}$<br>(kcal·mol<sup>-1</sup>) | $\Delta G_{\text{KL,blunt}}$<br>(kcal·mol<sup>-1</sup>) | $\Delta G_{\text{KL,blunt,shift}}$<br>(kcal·mol<sup>-1</sup>) |
| --- | --- | --- | --- | --- | --- |
| AA/TT | -1.00 | -1.11 | 0.11 | -1.36 | -0.79 |
| AT/AT | -0.88 | -1.34 | 0.46 | -2.35 | -1.78 |
| TA/TA | -0.58 | -0.19 | -0.39 | -1.01 | -0.44 |
| GG/CC | -1.84 | -1.44 | -0.40 | -1.64 | -1.07 |
| GC/GC | -2.24 | -2.17 | -0.07 | -3.42 | -2.85 |
| CG/CG | -2.17 | -0.91 | -1.26 | -2.06 | -1.49 |
| TG/CA | -1.45 | -0.55 | -0.90 | -0.81 | -0.24 |
| AG/CT | -1.28 | -1.06 | -0.22 | -1.60 | -1.03 |
| TC/GA | -1.30 | -1.43 | 0.13 | -1.39 | -0.82 |
| GT/AC | -1.44 | -1.81 | 0.37 | -2.03 | -1.46 |

First column indicates the base pair step, $\Delta G_{\text{KL,SantaLucia}}$ corresponds to the consensus nearest neighbor parameters of dsDNA stability of SantaLucia <sup>11</sup>, $\Delta G_{\text{KL,nick}}$ indicates stacking free energies extracted from dsDNA nicking experiments <sup>32</sup>, the free energy difference in column 4 corresponds to the expected base pairing contribution. $\Delta G_{\text{KL,blunt}}$ corresponds to stacking free energies of dsDNA blunt ends obtained from single molecule experiments <sup>35</sup>, The last column indicates $\Delta G_{\text{KL,blunt}}$ shifted by an offset of 0.57 kcal·mol<sup>-1</sup> (to obtain the same offset as for the $\Delta G_{\text{KL,nick}}$ )

**Table 2.** Base pairing contribution of each base pair step

| Base pair step<br>5'-3'/5'-3' | $\Delta G_{BP,1}$<br>(kcal·mol<sup>-1</sup>) | $\Delta G_{H-bond,1}$<br>(kcal·mol<sup>-1</sup>) | $\Delta G_{BP,2}$<br>(kcal·mol<sup>-1</sup>) | $\Delta G_{H-bond,2}$<br>(kcal·mol<sup>-1</sup>) |
| --- | --- | --- | --- | --- |
| AA/TT | -1.40 | -0.70 | -1.49 | -0.74 |
| AT/AT | -1.21 | -0.61 | -1.12 | -0.56 |
| TA/TA | -1.31 | -0.65 | -1.20 | -0.60 |
| GG/CC | -2.16 | -0.72 | -2.25 | -0.75 |
| GC/GC | -2.41 | -0.80 | -2.34 | -0.78 |
| CG/CG | -2.61 | -0.87 | -2.46 | -0.82 |
| TG/CA | -2.02 | -0.81 | -2.15 | -0.86 |
| AG/CT | -1.69 | -0.69 | -1.70 | -0.68 |
| TC/GA | -1.63 | -0.65 | -1.78 | -0.71 |
| GT/AC | -1.67 | -0.67 | -1.74 | -0.70 |
| Average contribution<br>per formed H-bond | - | -0.717+/-0.08 | - | -0.720+/-0.09 |

The base pairing contributions, $\Delta G_{BP,1}$ , and free energies per formed hydrogen bond, $\Delta G_{H-bond,1}$ , were obtained using the nicking free energies to calculate the dinucleotide stacking equilibrium<sup>32</sup>. The corresponding data in the last 2 columns are based on stacking free energy contributions obtained by Kilchherr et al. (<sup>35</sup>). Free energies per formed H-bond were calculated by dividing the base pairing contributions for the first 3 rows by 2, rows 4-6 by 3 and last 4 rows by 2.5.

In the following we focus on the base pair step contributions and do not consider effects due to DNA termini. We shortly summarize the current model of partitioning of dsDNA stability into base pairing and stacking contributions. In order to partition the contribution of a base pair step KL (K and L stand for A, C, G or T) to the stability of DNA into stacking and base pairing contributions Frank-Kamenetskii and coworkers ^32^ use,

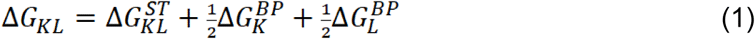

Where ΔG_KL_ is the total melting free energy parameter for a KL base pair step and the ΔG^BP^_K_ and the ΔG^BP^_L_ indicate the pairing contributions of base pair steps K and L, respectively. Since the DNA stability is calculated from overlapping base pair steps the factor ½ is necessary not to double count base pairing contributions. It is important to note that the stacking and base pairing contributions in this model represent effective contributions that take into account the nucleic acid backbone structure and flexibility. Hence, for example stacking does not represent here the stacking of isolated nucleobases but the effective stacking of bases in the context of a DNA backbone. In the model by Frank-Kamenetskii and coworkers for the stacking contribution ΔG^ST^_KL_ to double strand formation it is assumed that those are directly reflected by the free energies obtained from dsDNA nicking experiments (see Introduction section and reference ^32^). One should keep in mind, however, that nicking is a monomolecular conformational transition reaction whereas double strand formation involves two molecules and therefore the free energy is given with respect to a standard state (1 M) concentration of partners. A direct comparison usually involves an offset of the free energies (entropic contribution that depends on the chosen standard state). However, it turns out that the entropic contribution to the nicking process and to the process of adding a base pair to a dsDNA are very similar ^8^ and hence may justify the direct use of nicking free energies in eq. 1. With an appropriate choice for the base pairing contribution of an A:T and a G:C base pair it is then possible to obtain a linear dependence of the stability of long random DNA on the G/C content and to derive an independent set of nearest neighbor parameters. This is achieved using a predicted curve obtained using the consensus nearest neighbor model for the dsDNA stability as reference ^11^. The predicted base pairing contribution is slightly positive in case of A:T (∼0.09 kcal·mol^-1^) and close to zero for a G:C base pair (∼-0.1 kcal·mol^-1^) indicating a neglectable or slightly opposing contribution to dsDNA formation ^32^.

However, one would expect that the predicted base pairing contributions for A:T and G:C should be at least similar in all considered base pair steps. If one uses the consensus nearest neighbor model as reference and simply subtracts the effective ΔG^ST^_KL_ stacking contribution based on nicking experiments one should obtain the effective base pairing contribution (according to equation 1). Contrary to this expectation, the comparison gives very non-uniform base pairing contributions (see column 4 of Table 1). These can vary for A:T base pairs between negative (e.g. −0.39 kcal·mol^-1^ for a TA/TA step and positive contributions (0.46 kcal·mol^-1^ for an AT/AT step). In case of G:C pairing this variation is even larger (Table 1).

Secondly, if indeed the stacking of the DNA is the main driving force for dsDNA formation then the pre-stacking of the DNA (already in the unbound ssDNA) should destabilize dsDNA (because a gain of stacking upon dsDNA formation is then not anymore possible). Such prediction is counterintuitive and in order to qualitatively check this possibility we performed comparative free energy simulations on dissociating/forming a 2-base pair dsDNA from two ApT dinucleotides. The dsDNA molecules were dissociated using the Umbrella Sampling (US) technique (see Materials and Methods for details) and the associated free energy change was recorded (Figure 2). In one US simulations the dinucleotides were free to unstack during all stages of the simulation (black line in Figure 1). In the second US simulation unstacking was suppressed by an additional restraint (see Material and Methods, Supporting Information Table S1-3). The calculated free energy change was found to be significantly higher in case of suppressing the unstacking (stabilizing the bound form) which indicates that the gain in stacking upon dsDNA formation is not the driving force for the association. One may doubt that such free energy simulations can give a realistic quantitative insight into dsDNA stability but it may give at least a qualitative hint on the contributions. We proceed below by considering and re-analyzing only the experimentally measured contributions.

**Figure 1.**
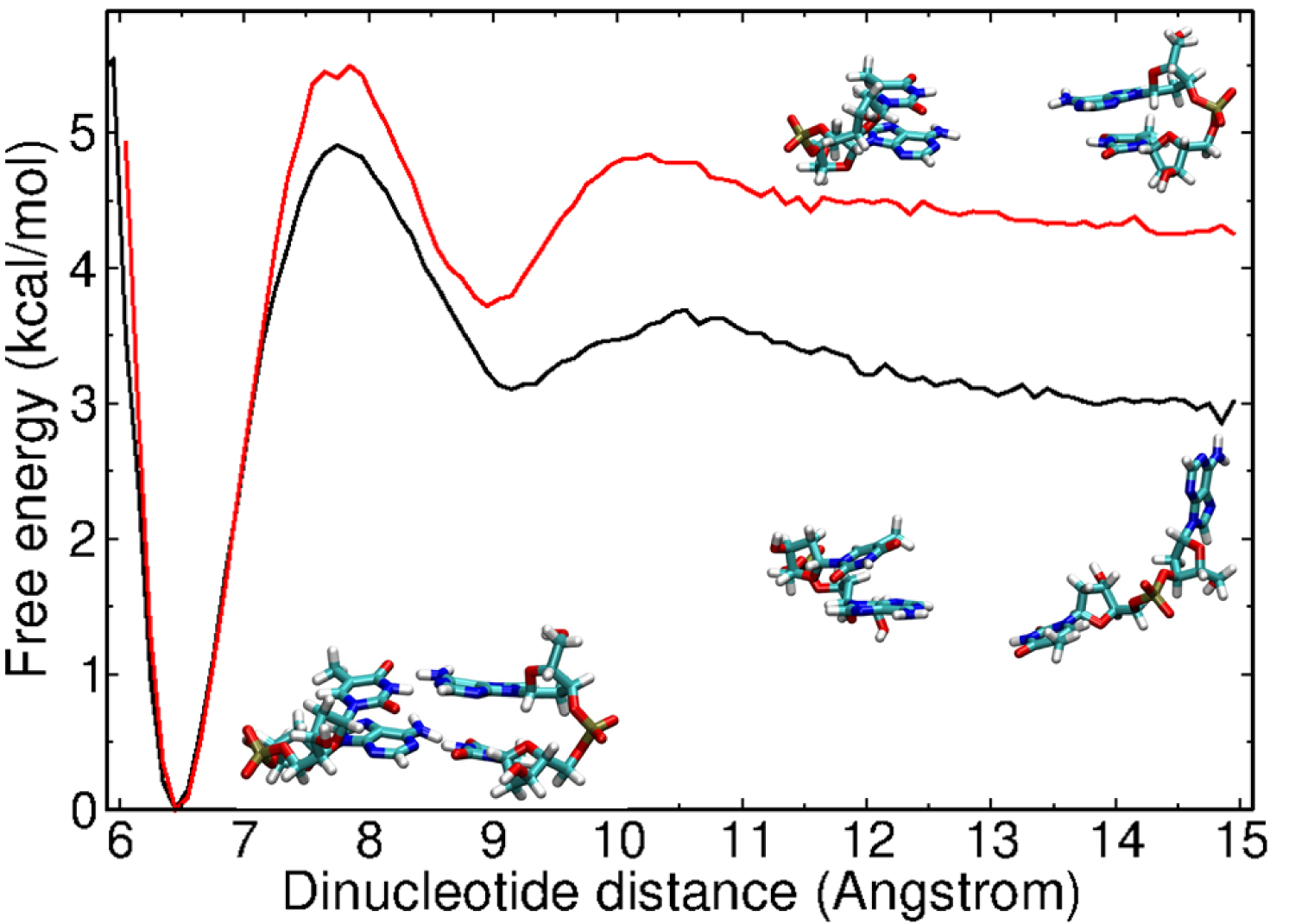
Calculated potential-of-mean force (PMF) or free energy change for the dissociation/association of two AT dinucleotides to form a dsDNA. The PMFs were calculated along a separation distance coordinate using the Umbrella Sampling (US) method. In one case the geometry of the AT stacking was kept near B-form geometry for the entire PMF calculation using appropriate restraints (red line) and in the other simulation the nucleobases were free to unstack during the simulation (black line). Snapshots of the base paired geometry (near the free energy minimum) and for the separate states (to the right of the panel) are indicated as stick models (the fully stacked separate state near the red line and a partially unstacked conformation in the unbound state below the black line).

**Figure 2.**
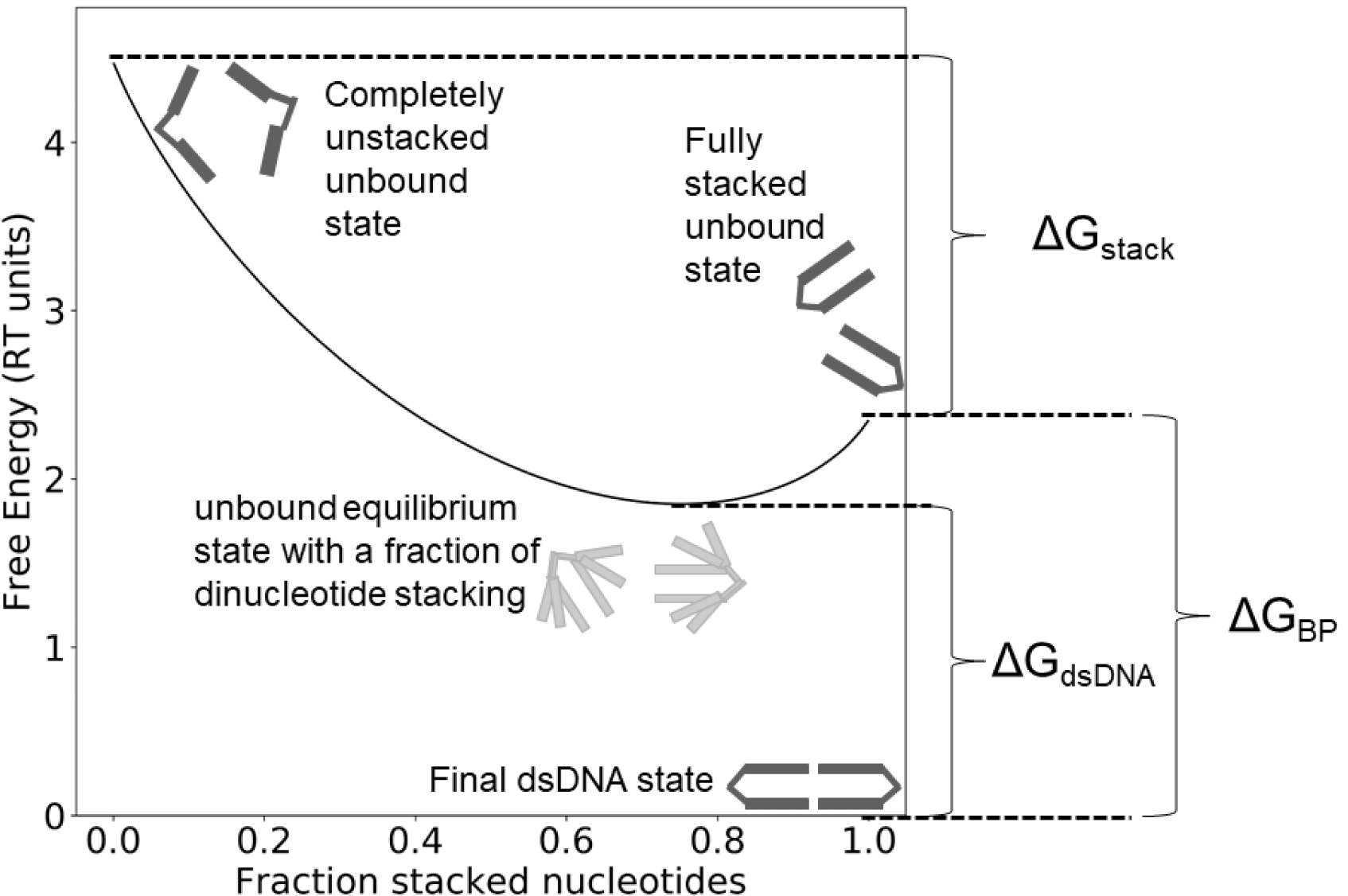
Schematically view of the free energy function (eq. 4) for describing the dsDNA formation process. The lowest free energy (zero) corresponds to a completely formed dsDNA base pair step. The free energy curve starts at two completely unstacked dinucleotides. At equilibrium (minimum free energy state) a fraction of these steps is in a stacked conformation (this fraction corresponds to the equilibrium constant determined by the experimentally measured stacking free energy, e.g. by nicking experiments). The minimum free energy state corresponds to the unbound reference state. In the diagram it is therefore placed at a free energy level that is shifted relative to the dsDNA level (zero free energy) by the consensus nearest neighbor free energy parameter for the corresponding base pair step. Upon full stacking of the dinucleotide steps a free energy level is reached that is lower than the completely unstacked state by the measured stacking free energy (indicated in the diagram). The final base pairing free energy must offset the cost for reaching the fully stacked state and must also account for the measured nearest neighbor stability contribution of the selected base pair step.

### An alternative model of dsDNA formation and stability

In eq. 1 for calculating the stability of dsDNA the nicking stacking free energies (Table 2) enter as stability parameters. These correspond to the transition from a fully unstacked configuration at the nick to a fully stacked state (like in B-dsDNA, resulting basically in the same electrophoretic mobility^32^). For the process of dsDNA formation this view considers the reference state (unbound state) as a fully unstacked ssDNA (the whole stacking energy enters into dsDNA formation). However, it is also possible to interpret the nicking free energies as driving force for forming stacked DNA conformations (like in B-DNA) already in the unbound single-stranded state. In this case the dsDNA formation process can be considered as transferring the partially stacked ssDNA to a fully stacked state (this actually costs free energy as we will see below) and forming the base pairing with the partner strand. The process and the associated free energy changes are illustrated in Figure 2. The nicking free energy for a given base pair step, e.g. AA/TT, can be interpreted as a contribution of A on A (in the first strand) and T on T (in the second strand). In case of symmetric steps (e.g. AT/AT) we have two identical stacking contributions (on both strands one has an A on T stack). We will see below that a splitting into contributions from symmetric or asymmetric steps plays no role for the further considerations. Note also, that both the nicking and the single strand stacking are monomolecular reactions so no free energy offsets need to be considered. For the change in free energy change (driving force) along the fraction of stacked conformations, x, we have,

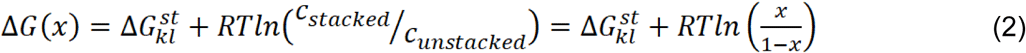

R indicates the gas constant and T the temperature. The ΔG^st^_kl_ now indicates the stacking free energy for a dinucleotide step (based on the nicking free energy), the ΔG(x) (free energy per fraction of stacking) is zero exactly when the fraction of the stacked states reaches the equilibrium (offsets ΔG^st^ _kl_). Hence, the free energy function along the reaction coordinate is given by (for convenience, we set RT=1),

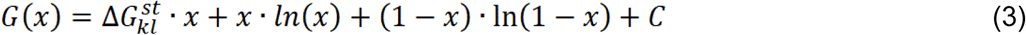

with C being an integration constant. This free energy function has its minimum exactly at the equilibrium fraction of stacked conformations (unbound state) and gives as difference between fully unstacked (x=0) and fully stacked states (x=1) exactly the measured ΔG^st^ _kl_. It is more useful than eq. (2) to illustrate the process of double strand formation and the associated free energy changes. To cover now the process of double strand formation of a full base pair step starting from completely unstacked dinucleotide partners we combine the G(x) for the two dinucleotides and substitute C such that we reach the experimental measured nearest neighbor base pair step free energy (with opposite sign) at the minimum of the curve. This is achieved by subtracting the minimum free energy of eq. (3), Gmin, and adding the nearest neighbor base pair step free energy (with opposite sign). For an AA/TT step, as an example, we get:

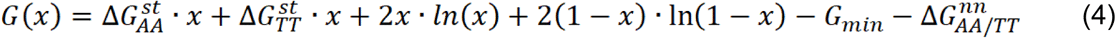

Note, that the sum of ΔG^st^_AA_ and ΔG^st^_TT_ represent in our treatment the measured nicking free energy for an AA/TT step. Note also, that the position of the free energy minimum represents the equilibrium stacking in the unbound state and its free energy difference relative to the base paired state is given by the nearest neighbor base pair step free energy (with opposite sign, illustrated in Figure 2).

Exactly this equilibrium position corresponds to the unbound reference state (and not the completely unstacked dinucleotides!). In the model the experimental measured free energy for forming a double-stranded base pair step (=nearest neighbor parameter) is the difference between free energy of the base pair step minus the free energy of the equilibrium partially stacked state of the two separate dinucleotide steps (illustrated in Figure 2). The plot in Figure 2 also shows that a free energy cost is associated with the formation of a fully stacked state of the dinucleotides (still unbound) relative to the equilibrium state. A convenient feature of the free energy plot in Figure 2 is that one can directly extract the free energy of base pair formation (that is the free energy at x=1, with the fully stacked dinucleotides relative to the double stranded base pair step, represented by a free energy level of zero in Figure 2). Depending on the stacking stability the equilibrium fraction of stacking and the penalty for reaching the fully stacked state will vary for each base pair step (see Figure 3). Also, for each case the equilibrium level of stacking in the unbound states of separate dinucleotides varies as expected.

**Figure 3.**
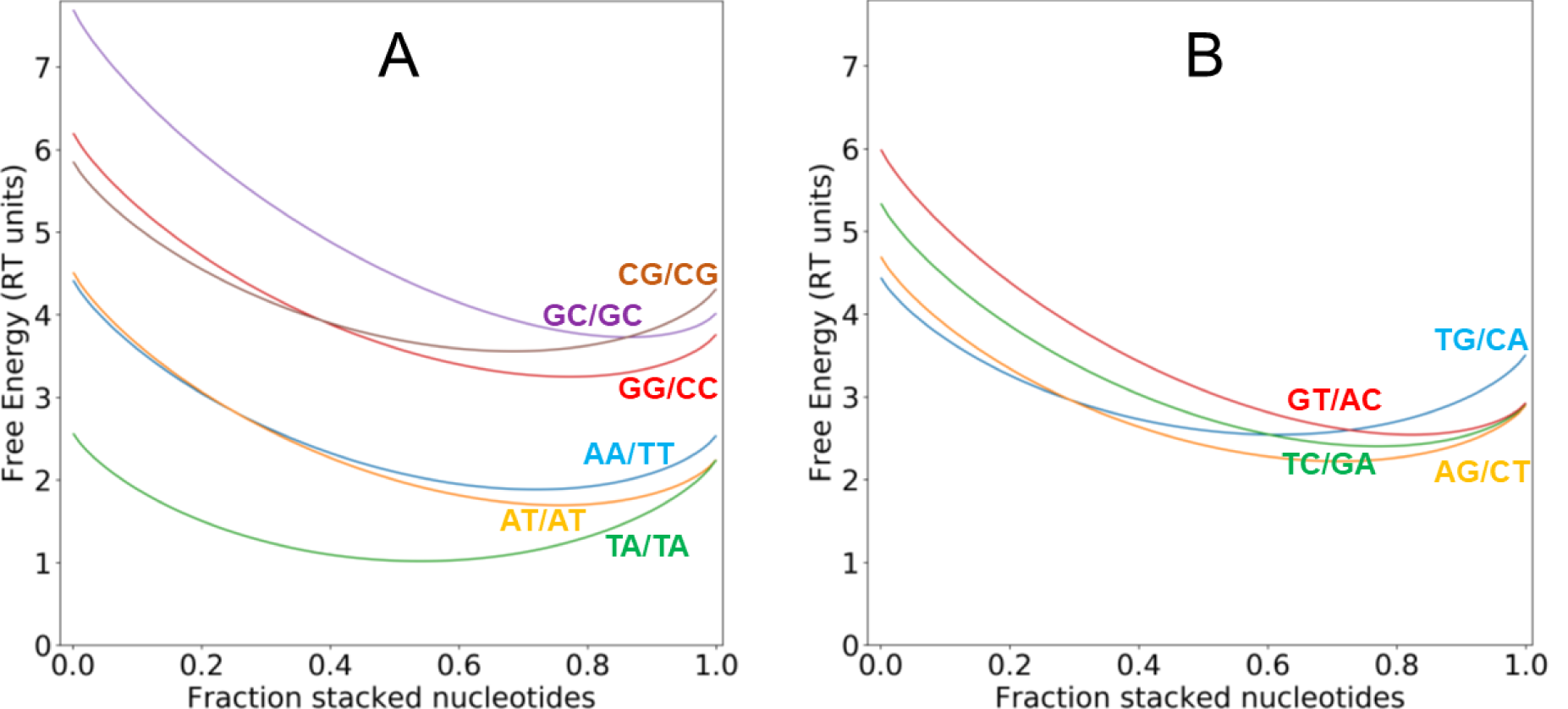
(A) Free energy vs. fraction of stacked dinucleotides for the AA/TT (blue), the AT/AT (orange), TA/TA (green), the GG/CC (red), GC/GC (violet) and CG/CG (brown) cases. (B) same as in (A) but for the “mixed” TG/CA (blue), AG/CT (orange), TC/GA (green) and GT/AC (red) base pair steps. The zero free energy level corresponds to the formed dsDNA state. The minimum free energy in each case corresponds to the nearest neighbor free energy contribution of the corresponding base pair step (for details on the scheme see legend of Figure 2).

The free energy curves of the steps that consist only of G:C base pairs consistently end up at higher free energy levels (at x=1) than those that contain only A:T pairs (Figure 3A). Also, the curves based on steps that involve only G:C base pairs reach almost the same free energy levels indicating that the base pairing contribution is very similar in the model. The same result is seen for the cases that only involve A:T base pairs. “Mixed” base pair steps (with one G:C and one A:T pair) reach intermediate free energies in between the pure G:C and A:T levels (Figure 3B). The average base pairing contribution follows a ratio of 3.3/2.6/2 for pure G/C, mixed and pure A/T steps (Table 2), respectively. This is very close to the expected ratio of 3/2.5/2 if one just considers the ratio of formed hydrogen bonds. However, one would expect that the base pairing contribution for steps with only G:C pairs such as GG/CC, GC or CG (or only A:T pairs) should be exactly the same (in terms of numbers of H-bonds) but the curves do not end up at exactly the same point for x=1. This is likely due to the limited accuracy of the experimental free energy determinations but also reflects small variations in hydrogen bonding stability due to the different local environments in each case (see below).

Recently, base pair stacking free energies were measured independently using a completely different methodology based on the DNA origami technique ^35^. Those base pairing free energies were obtained from the separation of two pairs of blunt end (double) dsDNA molecules and correspond to free energies measured with respect to a 1 M reference concentration of the partners. Hence, these free energies are comparable to the nicking free energies only with respect to an offset free energy (or only differences between two selected base pair steps can be directly compared). However, since the nicking free energies are monomolecular free energies we can “shift” the free energies from the origami studies to the same mean free energy (as the nicking free energies) (see Table 1, column 6). This effectively removes the offset. Using now instead of the nicking free energies the “corrected origami” stacking free energies ^35^ gives qualitatively very similar free energy curves vs. fraction of dinucleotide stacking (Supporting Information, Figure S2). In addition, quite similar mean base pairing free energy contributions are obtained (Table 2). This result further reassures the robustness of the theoretical model.

It is also possible to extract the free energy penalties to bring the dinucleotides into a B-type stacking in order to form a base pair (Table 3) from the curves shown in Figure 3 (and Supporting Information, Figure S2). The data in Table 3 illustrates that this penalty is for example quite high in case of a TA/TA step but small in case of a GC/GC step.

**Table 3.** Penalty for bringing dinucleotide partners into stacked conformation

| Base pair step<br>5'-3'/5'-3' | $\Delta G_{\text{stack\_penalty},1}$<br>(kcal·mol<sup>-1</sup>) | $\Delta G_{\text{stack\_penalty},2}$<br>(kcal·mol<sup>-1</sup>) |
| --- | --- | --- |
| AA/TT | 0.40<br>(0.147,0.253) | 0.495<br>(0.202,0.293) |
| AT/AT | 0.33<br>(0.164,0.164) | 0.24<br>(0.118,0.118) |
| TA/TA | 0.73<br>(0.363,0.363) | 0.62<br>(0.309,0.309) |
| GG/CC | 0.32<br>(0.105,0.218) | 0.41<br>(0.153,0.258) |
| GC/GC | 0.17<br>(0.087,0.087) | 0.10<br>(0.050,0.050) |
| CG/CG | 0.45<br>(0.224,0.224) | 0.29<br>(0.147,0.147) |
| TG/CA | 0.575<br>(0.287,0.287) | 0.70<br>(0.352,0.352) |
| AG/CT | 0.41<br>(0.154,0.260) | 0.42<br>(0.160,0.263) |
| TC/GA | 0.33<br>(0.219,0.106) | 0.485<br>(0.289,0.196) |
| GT/AC | 0.23<br>(0.115,0.115) | 0.30<br>(0.150,0.150) |

It is also of interest to investigate how accurate the stability of an arbitrary dsDNA sequence can be predicted based on assuming just a single base pairing contribution per H-bond (for every base pair) compared to the original nearest-neighbor model. The near-neighbor (NN) free energy of the following sequence segment (we assume it is a segment of a long sequence and neglect boundary contributions),

..GCTACAAG..

can be calculated as the sum of NN contributions (Table 1, column 2):

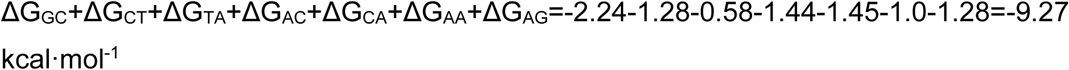

Using the current model with a unique mean pairing contribution per formed H-bond (−0.72 kcal·mol^-1^) we need to add the penalties for each step (Table 3, column 2) and add the number of formed H-bonds:

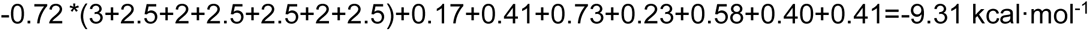

Hence, for this example the agreement is close to perfect. The application to 200 random sequences (of length 5 to 35) using a mean base pairing contribution per H-bond gives a standard deviation from the original nearest neighbor model of 0.78 kcal·mol^-1^ (Figure 4A) which is smaller than the mean prediction accuracy of the NN model^5^. Inspection of Table 2 indicates that the contribution per formed H-bond seems to be slightly larger for steps that contain only G:C base pairs (mean: −0.80 kcal·mol^-1^) vs. those containing only A:T pairs (mean: −0.65 kcal·mol^-1^) or mixed case (mean: −0.71 kcal·mol^-1^). Using these mean per H-bond contributions the standard deviation for the 200 random cases drops to 0.66 kcal·mol^-1^ (Figure 4B).

**Figure 4.**
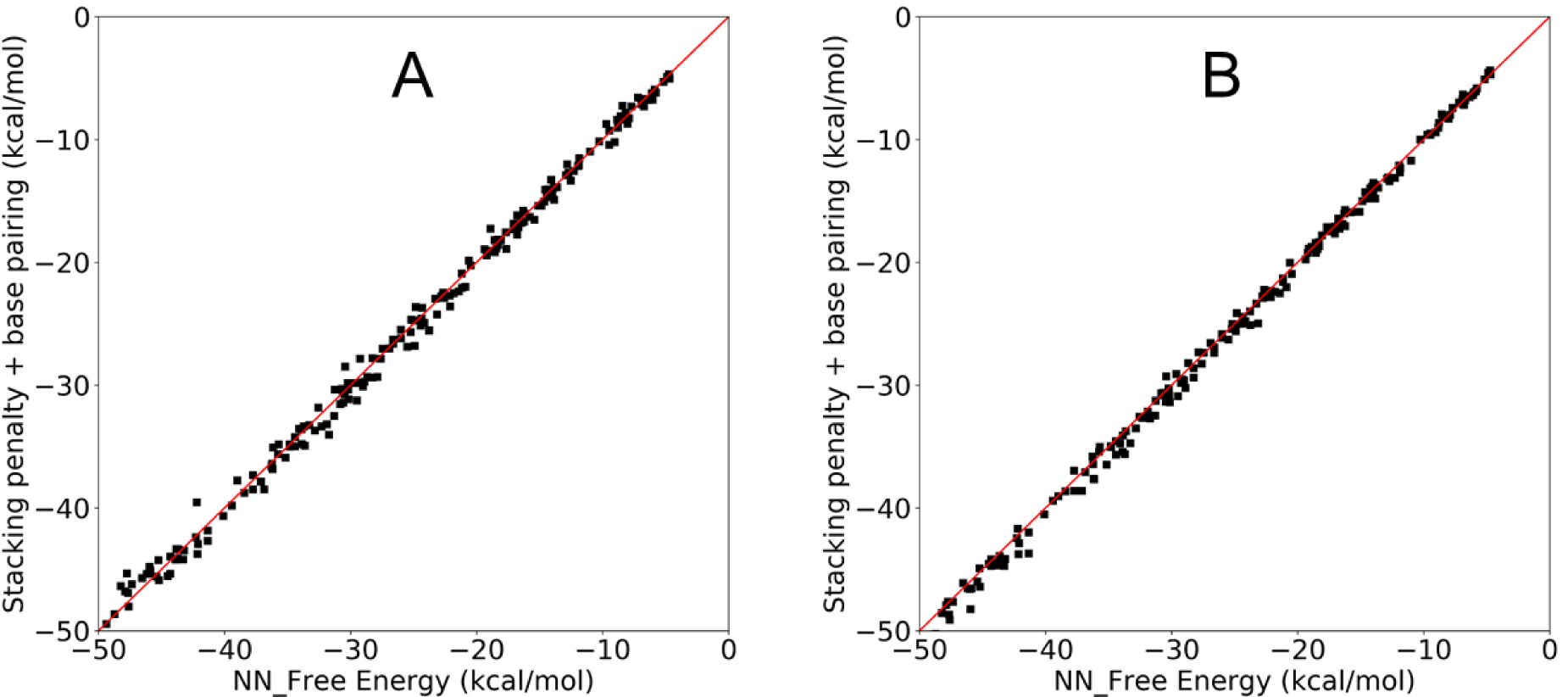
(A) Correlation of calculated dsDNA stabilities for 200 random sequences (length distribution 5-35) using the united nearest-neighbor model^11^ (x-axis) vs. the model employing a single effective free energy per formed H-bond (−0.72 kcal·mol^-1^) combined with free energy penalties to bring dinucleotides in a stacked B-DNA like conformation (extracted from Table 3). (B) same as (A) but using 3 different free energy contributions per formed H-bond (−0.65 kcal·mol^-1^ in of steps that involve only A:T pairing, −0.80 kcal·mol^-1^ in case of steps involving only G:C pairing and −0.71 kcal·mol^-1^in case of mixed steps). The red line represents a 1:1 correspondence.

Hence, based on the present re-analysis it is possible to assign a mean free energy per hydrogen bond formed during dsDNA formation of −0.72+/-0.08 kcal mol^-1^ that is consistent with all data (Table 2) and only narrowly distributed. Again reassuring, basically the same numbers are obtained if stacking data based on nicking or blunt end single molecule manipulation are used (Table 2, last row).

## Conclusions

A quantitative understanding of the stability of double stranded DNA is of fundamental importance for almost all research areas on nucleic acids. Both stacking as well as base pairing have been identified to contribute to dsDNA stability. Here, effective stacking and base pairing contributions are always considered, that means in the context of a nucleic acid backbone structure (taking into account the flexibility of the backbone that may energetically or entropically oppose or promote stacking and/or pairing). In recent experimental studies but also in several theoretical studies the importance of stacking contributions has been emphasized. Efforts to separate stacking and base pairing contributions lead to the view that stacking is the main component that drives dsDNA formation whereas base pairing forces make only a minor or even slightly opposing contribution^8,32^. However, such model assumes that in the unbound separate states of the two DNA strands the bases are unstacked and the “full” stacking free energy enters into the dsDNA formation process. This view is equivalent to considering for the association of two proteins the fully unfolded conformation of the two partners as the unbound configuration (unbound reference state). Hence, if we assume the two proteins are unfolded (“unstacked”) in the unbound state and only fold (“stack”) upon binding then the binding energy must be so strong that it can “pay the price” to fold the partner molecules. Hence, the folding free energy of the partners cannot be added to the binding free energy (it effectively opposes binding). For the dsDNA case if stacking is indeed the only driving force for double strand formation than bringing the partners already in a “pre” -stacked conformation should lower the binding affinity (between the two ssDNAs). Base pair step free energy simulations on an example in the present study indicate that this is unlikely being the case. Furthermore, just subtracting measured stacking free energies from experimental nearest neighbor base pair formation free energies gives widely varying base pairing contributions for steps that have the same number of A:T of G:C base pairs.

The present theoretical model of the stability of double stranded DNA is based on the assumption of a significant fraction of stacked single stranded nucleobases that depends significantly on the nucleotide sequence. Since the dsDNA formation process requires the transition of each dinucleotide partner to adopt a fully stacked (B-form-like) structure it predicts a (dinucleotide specific) penalty that needs to be overcome by the base pairing contribution (see Figure 2 and Table 3, corresponds to the difference ΔG_BP_-ΔG_dsDNA_). Hence, the final transition to a fully stacked state actually opposes the double strand DNA formation (depending on the sequence of the step). The final dsDNA base pairing needs to overcome this penalty and to contribute the nearest neighbor free energy increment. Contrary to previous models, base pairing makes therefore the decisive contribution to dsDNA stability. The model results in a physically consistent picture of the dsDNA formation process and allows one (for the first time) to extract an effective free energy per formed hydrogen bond in the dsDNA based entirely on experimental data. It should be emphasized that this is not the “energy” (or free energy) of any isolated H-bond but an effective free energy per formed H-bond under standard state conditions upon forming a dsDNA. It may also have significant implications for interpreting the stability of non-canonical motifs in DNA.

## Acknowledgements

I like to thank K. Liebl and H. Dietz for helpful discussions. This work was supported by the Deutsche Forschungsgemeinschaft (DFG) grant Za153/28-1.

## Supporting Information

**Figure S1.**
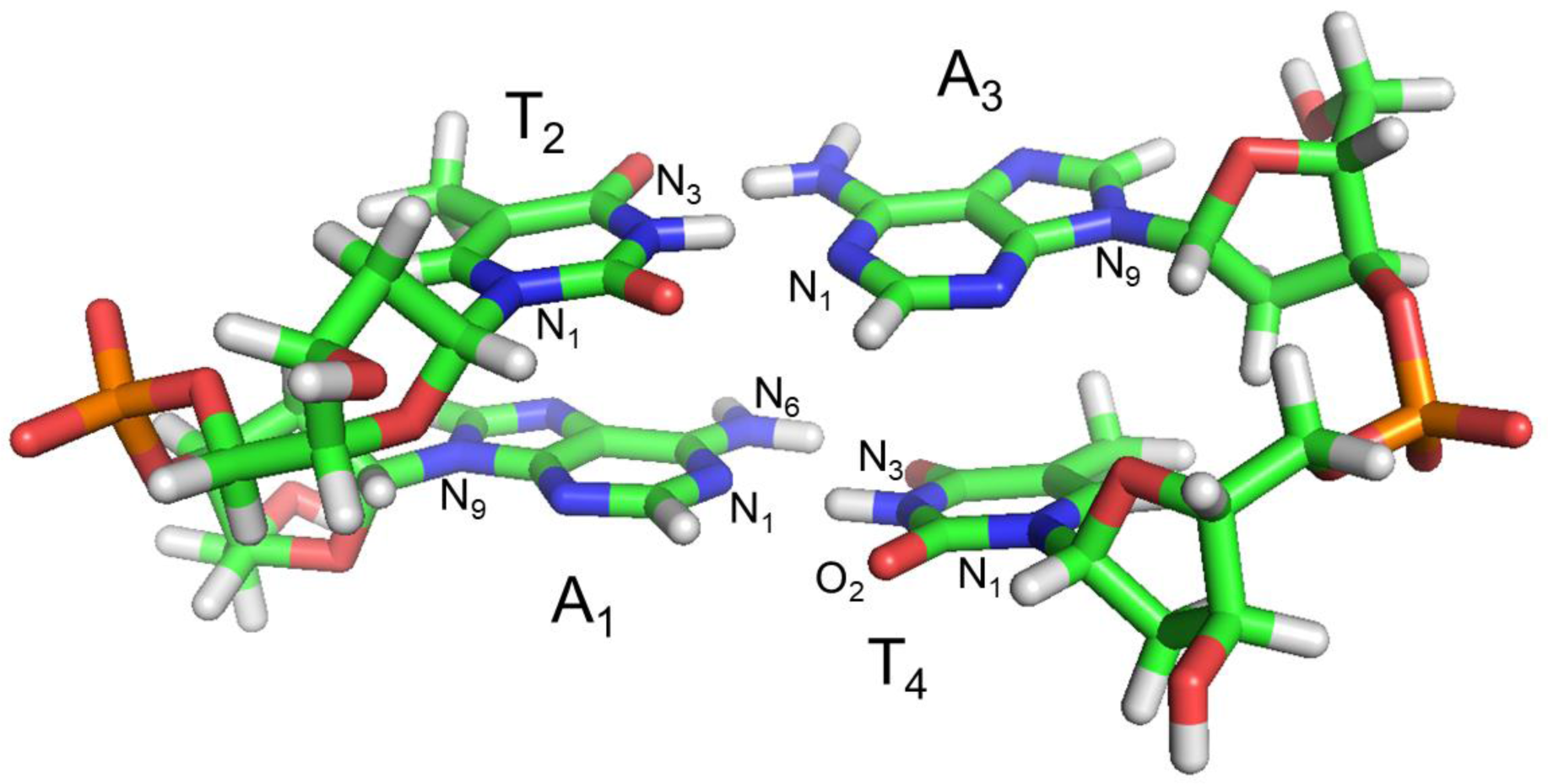
Illustration of a B-DNA base pair (atom-color-coded stick model) formed by the association of 2 AT dinucleotides. The nucleotides are marked by large letters and atoms relevant for the definition of the distance, angular and dihedral restraints used during the Umbrella Sampling simulations are marked by small letters.

**Table S1.** Definition of the center-of-mass distance reaction coordinate used in Umbrella ampling simulations to calculate the PMF for base pair formation.

| Equilibrium distance<sup>a</sup> in<br>quadratic potential | Force constant (kcal·mol<sup>-1</sup>·Å<sup>-2</sup>) |
| --- | --- |
| 6.0 Å | 3.0 |
| 6.5 Å | 3.0 |
| 7.0 Å | 3.0 |
| 7.5 Å | 3.0 |
| 8.0 Å | 2.5 |
| 9.0 Å | 2.0 |
| 10.0 Å | 2.0 |
| 11.0 Å | 1.5 |
| 12.5 Å | 1.0 |
| 14.0 Å | 1.0 |

<sup>a</sup>) The center-of-mass distance was calculated between all non-hydrogen atoms of the first nucleobase (A<sub>1</sub>) and the complementary nucleobase (T<sub>4</sub>) on the opposite strand.

**Table S2.** Angle and Dihedral restraints to determine the direction of dinucleotide association/dissociation during US simulations.

| Set of atoms defining the angular or dihedral restrain <sup>a</sup> | Equilibrium angle or dihedral angle in quadratic potential (degrees) | Force constant (kcal·mol<sup>-1</sup>·rad<sup>-2</sup>) |
| --- | --- | --- |
| A<sub>1</sub>(N<sub>6</sub>), A<sub>1</sub>(N<sub>1</sub>), T<sub>4</sub>(N<sub>3</sub>), T<sub>4</sub>(O<sub>2</sub>) | 180.0 | 25.0 |
| A<sub>1</sub>(N<sub>6</sub>), A<sub>1</sub>(N<sub>1</sub>), T<sub>4</sub>(N<sub>3</sub>) | 90.0 | 25.0 |
| A<sub>1</sub>(N<sub>1</sub>), T<sub>4</sub>(N<sub>3</sub>), T<sub>4</sub>(O<sub>2</sub>) | 90.0 | 25.0 |
| A<sub>1</sub>(N<sub>9</sub>), T<sub>4</sub>(N<sub>3</sub>), T<sub>4</sub>(N<sub>1</sub>) | 170.0 | 10.0 |
| A<sub>1</sub>(N<sub>9</sub>), A<sub>1</sub>(N<sub>1</sub>), T<sub>4</sub>(N<sub>1</sub>) | 170.0 | 10.0 |

<sup>a</sup>) The equilibrium angles were obtained from short unrestrained MD simulations of the dinucleotides in the base paired geometry. The force constants represent approximately the flexibility observed during these unrestrained simulations. **Note, the restraints of Table S2 were employed in all US simulations.**

**Table S3.** Additional Angle and Dihedral restraints to keep the dinucleotide states in a stacked state during all US simulations.

| Set of atoms defining the angular or dihedral restrain <sup>a</sup> | Equilibrium angle or dihedral angle in quadratic potential (degrees) | Force constant (kcal·mol<sup>-1</sup>·rad<sup>-2</sup>) |
| --- | --- | --- |
| A<sub>1</sub>(N<sub>1</sub>), A<sub>1</sub>(N<sub>9</sub>), T<sub>2</sub>(N<sub>1</sub>), T<sub>2</sub>(N<sub>3</sub>) | 39.0 | 25.0 |
| A<sub>3</sub>(N<sub>1</sub>), A<sub>3</sub>(N<sub>9</sub>), T<sub>4</sub>(N<sub>1</sub>), T<sub>4</sub>(N<sub>3</sub>) | 46.0 | 25.0 |
| T<sub>2</sub>(N<sub>1</sub>), A<sub>1</sub>(N<sub>9</sub>), A<sub>1</sub>(N<sub>1</sub>) | 70.0 | 25.0 |
| A<sub>1</sub>(N<sub>9</sub>), T<sub>2</sub>(N<sub>1</sub>), T<sub>2</sub>(N<sub>3</sub>) | 88.0 | 25.0 |
| A<sub>3</sub>(N<sub>9</sub>), T<sub>4</sub>(N<sub>1</sub>), T<sub>4</sub>(N<sub>3</sub>) | 86.0 | 25.0 |
| T<sub>2</sub>(N<sub>1</sub>), A<sub>1</sub>(N<sub>9</sub>), A<sub>1</sub>(N<sub>1</sub>) | 73.0 | 25.0 |

<sup>a</sup>) The equilibrium angles used as restraining reference values were obtained from short unrestrained MD simulations of the dinucleotides in the base paired geometry. The force constants represent approximately the flexibility observed during these unrestrained simulations. **Note, the restraints of Table S3 were only employed in US simulations to keep the dinucleotides stacked during simulations.**

**Figure S2.**
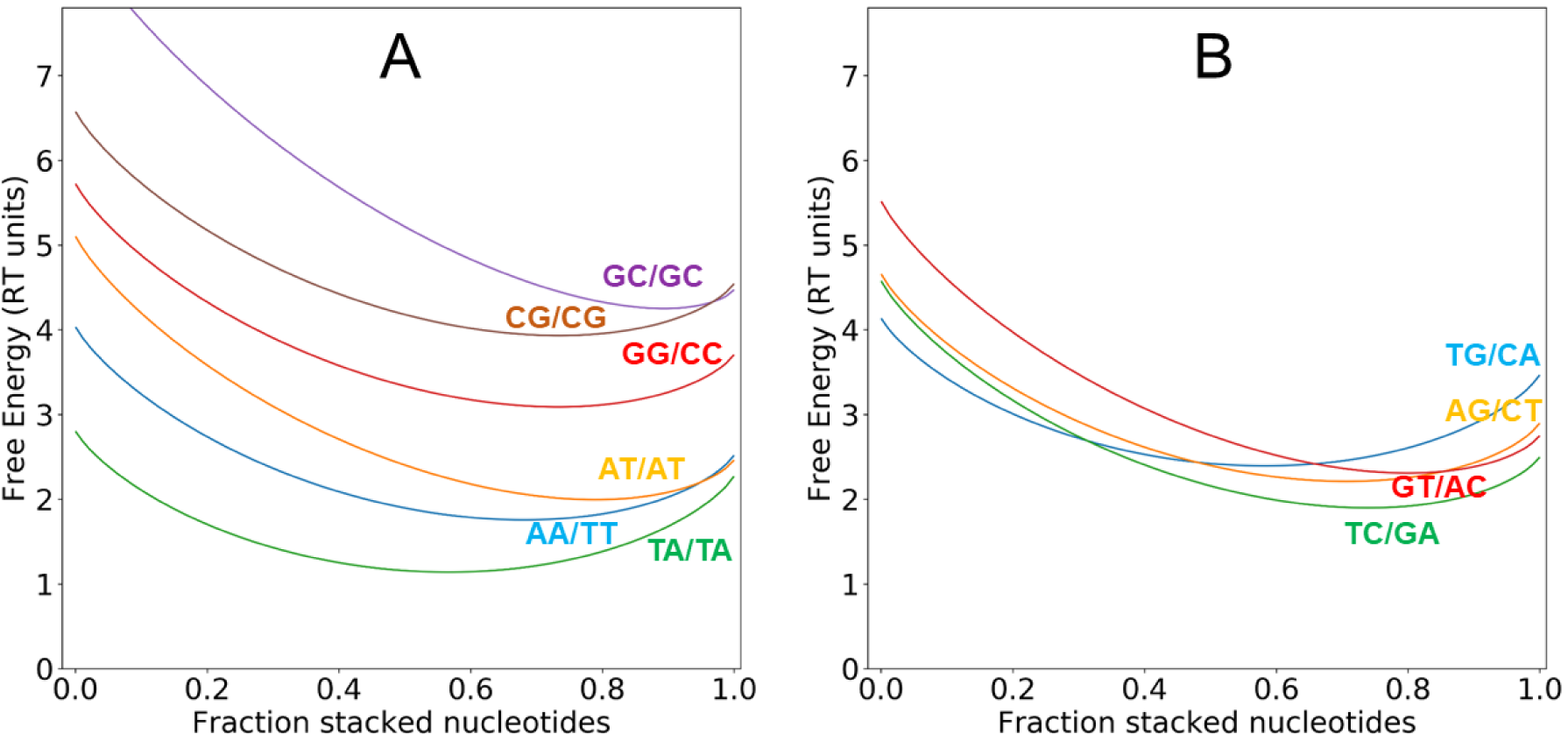
(A) Free energy vs. fraction of stacked dinucleotides for the AA/TT (blue), the AT/AT (orange), TA/TA (green), the GG/CC (red), GC/GC (violet) and CG/CG (brown) cases. (B) same as in (A) but for the “mixed” TG/CA (blue), AG/CT (orange), TC/GA (green) and GT/AC (red) base pair steps. The zero free energy level corresponds to the formed dsDNA state. The minimum free energy in each case corresponds to the nearest neighbor free energy contribution of the corresponding base pair step (for details on the scheme see legend of Figure 2 in the main text). In contrast to Figure 3 in the main manuscript the stacking free energies of Kilchherr et al. (1) were used.

